# Discovery of a new path for red blood cell generation in the mouse embryo

**DOI:** 10.1101/309054

**Authors:** Irina Pinheiro, Özge Vargel Bölükbaşi, Kerstin Ganter, Laura A. Sabou, Vick Key Tew, Giulia Bolasco, Maya Shvartsman, Polina V. Pavlovich, Andreas Buness, Christina Nikolakopoulou, Isabelle Bergiers, Valerie Kouskoff, Georges Lacaud, Christophe Lancrin

## Abstract

Erythropoiesis occurs through several waves during embryonic development. Although the source of the primitive wave is well characterized, the origin of erythrocytes later in embryogenesis is less clear due to overlaps between the different erythroid waves. Using the miR144/451-GFP mouse model to track cells expressing the erythroid microRNAs miR144/451, we identified cells co-expressing VE-Cadherin and GFP in the yolk sac between E9.5 and E12. This suggested the existence of hemogenic endothelial cells committed to erythropoiesis (Ery-HEC). We showed that these cells were capable of generating erythrocytes *ex vivo* and we demonstrated that the formation of Ery-HEC was independent of the *Runx1* gene expression. Using transcriptome analysis, we demonstrated that these cells coexpressed endothelial and erythroid genes such as *Hbb-bh1* and *Gata1* but we were surprised to detect the primitive erythroid genes *Aqp3* and *Aqp8* suggesting the formation of primitive erythrocytes at a much later time point than initially thought. Finally, we showed that enforced expression of *Gata1* in endothelial cells was enough to initiate the erythroid transcriptional program.

## Introduction

Hematopoiesis is thought to develop in the mouse embryo through three distinct waves, characterized by the appearance of distinct hematopoietic progenitors at various time points (Yoder, 2014). The first wave starts at E7 with the generation of primitive erythroid progenitors, some macrophages and megakaryocytes in the yolk sac (YS). These are still detected at E8.25, while at around E9 and E10.5 only some primitive macrophages persist. At E8.25 the second wave starts within the YS hemogenic endothelium (an endothelium with hematopoietic capacity), giving rise to erythromyeloid progenitors (EMPs). These are still detected at E9 and 10.5 of embryonic development, along with B and T lymphoid cells. Finally, the third wave is marked by the appearance of the first hematopoietic stem cells (HSCs), generated by the hemogenic endothelium in the aorta-gonad-mesonephros (AGM) region, around E10.5 (Yoder, 2014).

Erythroid cells are generated by all of the aforementioned hematopoietic waves, supplying the rapidly growing embryo with oxygen and playing a role in vascular remodeling. The generation of the first erythrocyte progenitors (Erythroid colony forming cells: EryP-CFC) is transient, as they are no longer detected at E9.5 (Palis et al., 1999, Isern et al., 2011). Nonetheless, erythrocytes continue to mature in a semi synchronous cohort while circulating in the bloodstream (Fraser et al., 2007; Kingsley et al., 2004). EMPs provide the growing organism with the second erythroid wave, prior and independently of the first HSCs emergence (McGrath et al., 2011). They start colonizing the fetal liver at E10.5 and are still detectable in the YS until E11.5 (Palis et al., 1999). Finally, a permanent third wave of erythroid cells is derived from HSCs that emerge from the AGM region at E10.5.

The overlap between distinct waves in the embryo as they enter circulation complicates the distinction between cell types. Nonetheless, the first erythroid wave can be distinguished from the rest due to their large nucleated cells and embryonic hemoglobin expression (mainly βH1 and εγ) (Palis, 2014). However, once they enucleate, primitive erythroid cells are harder to distinguish, especially from erythroid derived EMPs as they also express βH1 globin in low amounts (McGrath et al., 2011). Additionally, primitive erythroid cells also show expression of adult globins (β1 and β2) even though at lower levels (Palis, 2014). The first primitive erythroid progenitors were thought to arise from a mesodermal precursor (Ema et al., 2006; Isern et al., 2011). On the other hand, EMPs and blood progenitor/stem cells generated in the later waves arise from mature endothelial cells with hemogenic potential, through a process termed Endothelial to Hematopoietic Transition (EHT) (Lancrin et al., 2009). Interestingly, Stefanska et al have shown recently that primitive erythroid cells are generated in the yolk sac from hemogenic angioblast cells between E7.5 and E8.0 corresponding to the establishment of the capillary plexus (Garcia et al., 2014), when the vasculature is still immature (Stefanska et al., 2017).

The transcription factor RUNX1 plays a crucial role during EHT, being responsible for the loss of endothelial properties during blood generation (Lancrin et al., 2009, Lancrin et al., 2012, Thambyrajah et al., 2016). Even though RUNX1 is dispensable for the generation of the first primitive erythroid cells, it plays a role in the maintenance of their morphology and correct gene expression (Yokomizo et al., 2008). Additionally, the loss of RUNX1 is detrimental for the generation of EMPs and definitive hematopoietic cells derived from the hemogenic endothelium (Chen et al., 2009). On the other hand, the exact role of the transcription factor GATA1 in EHT remains relatively unknown although it is clearly critical for erythroid cell formation in all three waves of blood cell formation.

In this study, we asked whether we could find in the yolk sac hemogenic endothelial cells specifically generating erythroid cells after E9.5; i.d. following vascular remodeling and production of a highly structured vascular network (Garcia et al., 2014). To investigate this question, we used a previously described mouse model, in which GFP is used to label the transcription of the erythroid specific microRNAs miR144/451, which are expressed at all stages of erythroid differentiation (Rasmussen and O’Carroll, 2011). We found a novel endothelial population that displays high expression of primitive erythroid genes, possibly representing a hemogenic endothelium subset prone to give rise to erythroid cells only. We suggest for the first time that primitive erythroid cells can be generated at later time points than previously described, after the onset of both primitive and definitive erythroid waves. In addition, we showed that this population can emerge in the absence of *Runx1* and that the overexpression of *Gata1* specifically in endothelium, could initiate the erythroid transcriptional program.

## Results

### A new cell population with endothelial and erythroid gene expression is found in the yolk sac between E9.5 and E12

We used a miR144/451^+/GFP^ mouse model, in which the erythroid lineage can be tracked by GFP expression (Rasmussen et al., 2011) to find out if an endothelial population with erythroid differentiation capacity exists between E9.5 and E12, i.e. after the primitive hematopoiesis wave. Yolk Sacs (YS) were isolated at different time points, dissociated and cells were subsequently analyzed for co-expression of VE-Cad (endothelial marker) and miR144/451-GFP by FACS (Fig. 1A and B). A population that expresses both GFP and the cell surface marker VE-Cad was found at E9.5. The percentage of VE-Cad^+^GFP^+^ increases slightly from E9.5 (0.0116%), reaching its peak at E11.5 (0.0378%), before reducing slightly at E12. This population represents approximately 0.1% of all yolk sac VE-Cad^+^ cells at all tested times. However since there is an increase in the total cell numbers in the YS as the embryo develops, there is a net increase of double positive cells during development (Fig. 1C). As shown in Fig. 1C, a small number of double positive cells are found at E9.5 (≈21 cells), increasing significantly at E12 to an average of 425 cells in the YS.

**Figure 1:**
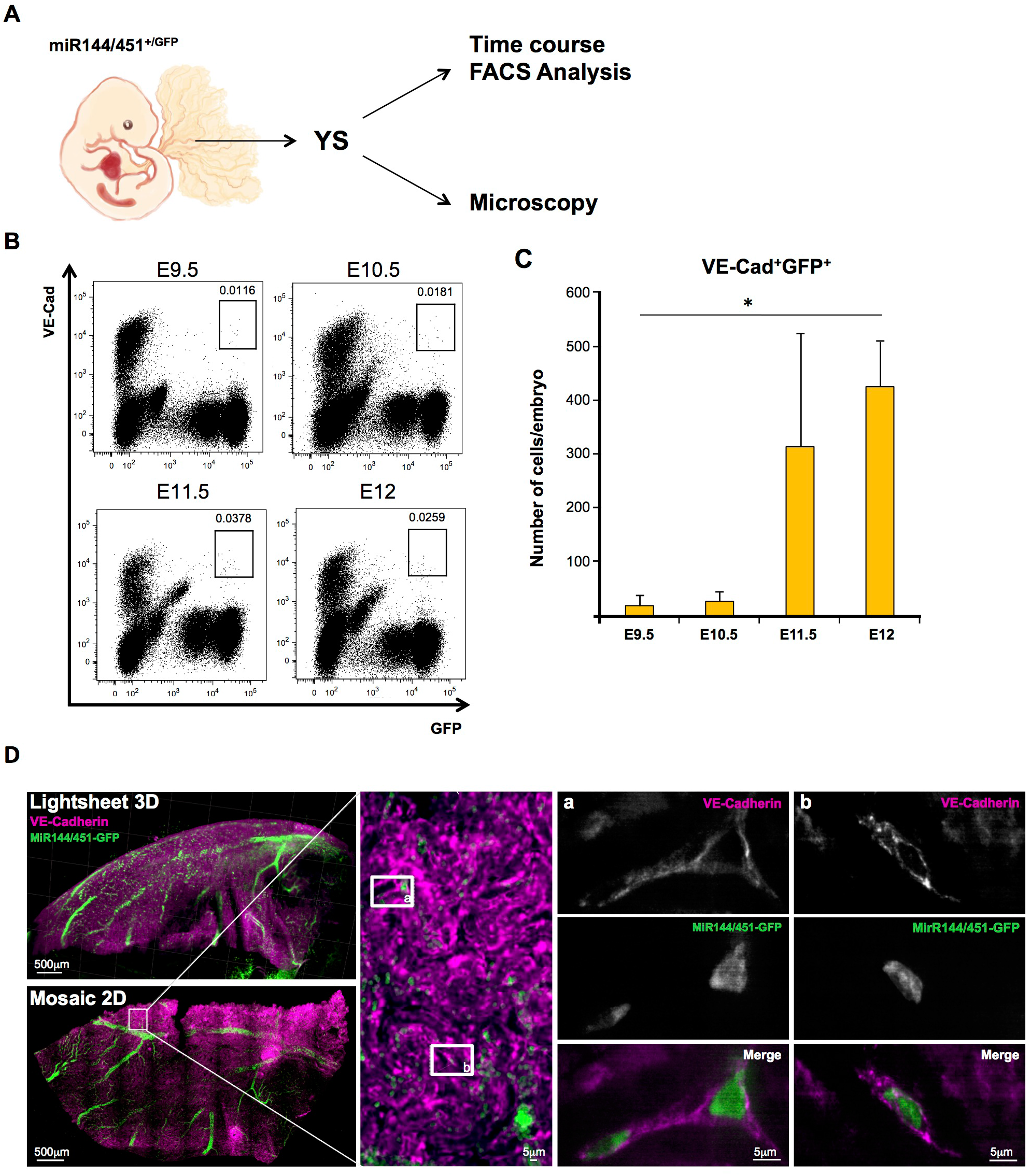
Identification of new population of endothelial cells in the yolk sac. (A) Schematic representation of the experimental protocol for FACS analysis and microscopy of miR144/451^+/GFP^ YS VE-Cad^+^GFP^+^ population, at different time-points. (B) FACS plots showing VE-Cad and GFP expression in YS at different time points during embryo development. (C) Bar graphs representing the average number ± SD of VE-Cad^+^GFP^+^ cells per embryo YS, at distinct time points. N=3 independent experiments. Kruskal-Wallis test, *p<0.05. (D) Immunofluorescence analysis of miR144/451^+/GFP^ YS. On the left panels, low-magnification views of the tissue by lightsheet 3D and Mosaic 2D are shown. A small area in the mosaic 2D was used for high magnification microscopy. Two areas labeled “a” and “b” containing cells coexpressing VE-Cad and GFP were found and further analyzed (right panels).

We then used the miR144/451^+/GFP^ mouse model to visualize the VE-Cad^+^GFP^+^ cells in the YS by fluorescence microscopy at E11.5 and found cells co-expressing VE-Cadherin and GFP (Fig. 1D). These cells appear to be flat and elongated, thus exhibiting a morphology that is similar to endothelial cells.

### Single cell transcriptome analysis demonstrated that miR144/451^+^ endothelial cells expressed the erythroid genes *Gata1* and *Hbb-bh1*

The molecular signature of this newly discovered population was assessed by singlecell q-RT-PCR between E10.5 and E12 (Fig. 2A). We analyzed the gene expression of 95 genes, including endothelial, hematopoietic and erythroid specific markers (Table S1), at the single cell level for the following populations: VE-Cad^−^GFP^+^, VE-Cad^+^GFP^−^ and VE-Cad^+^GFP^+^.

**Figure 2:**
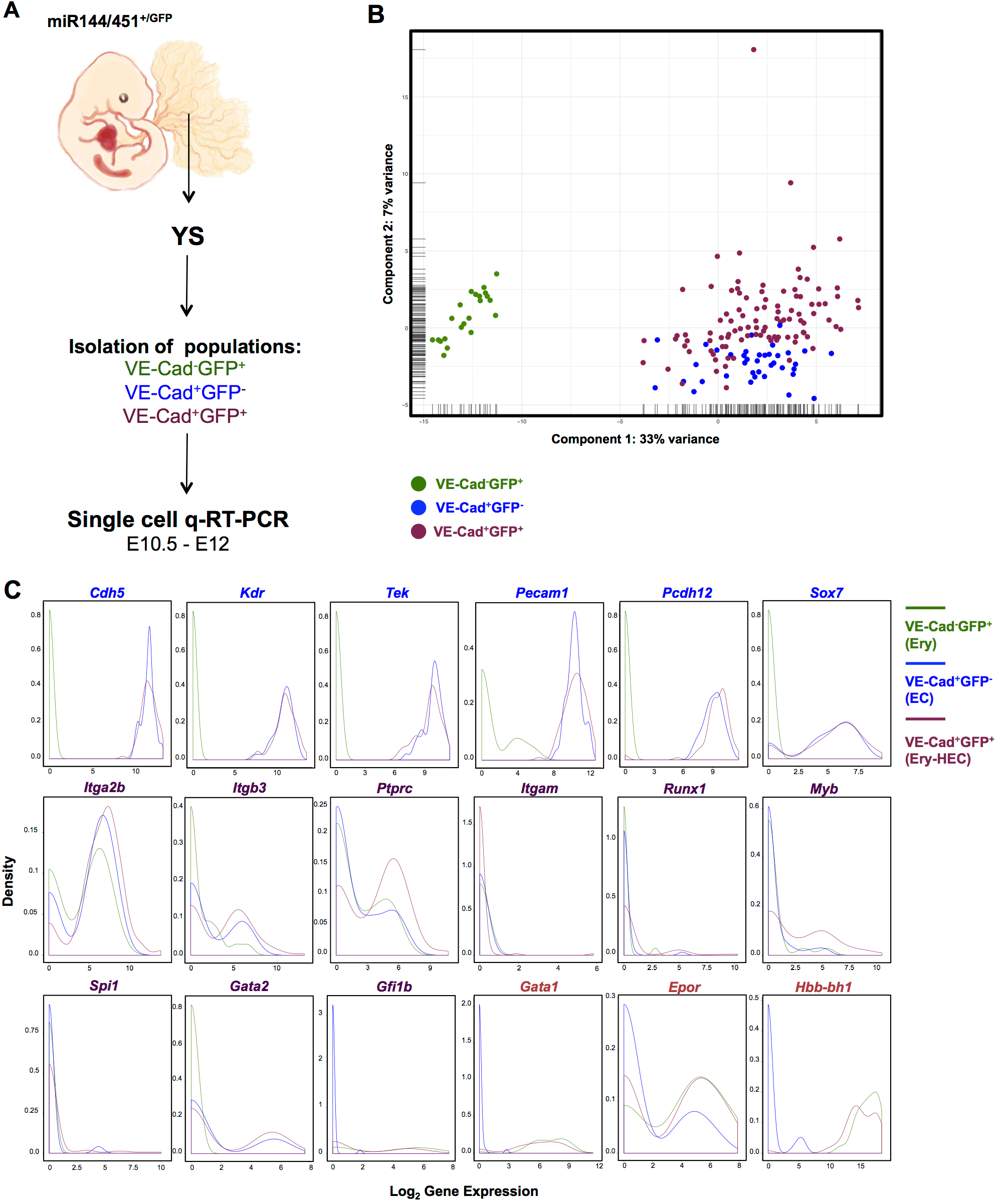
miR144/451^+^ endothelial cells express *Gata1, Epor* and *Hbb-bh1*. (A) Schematic representation of the experimental protocol for single cell q-RT-PCR of miR144/451^+/GFP^ YS VE-Cad^+^GFP^+^ populations, at different time-points. VE-Cad^−^GFP^+^ and VE-Cad^+^GFP^−^ cells were used as controls. (B) PCA plot showing the distribution of the single cells analyzed by single cell q-RT-PCR. (C) Density plots showing the gene expression distribution of selected genes: endothelial (*Cdh5, Kdr, Tek, Pecam1, Pcdh12, Sox7*), hematopoietic (*Itga2b, Itgb3, Ptprc, Itgam, Runx1, Myb*) and erythroid (*Gata1, Epor, Hbb-bh1*). The colored lines indicate the different populations. See also Figure S1.

Following single cell q-RT-PCR analysis, Principal Component Analysis (PCA) was performed. The VE-Cad^−^GFP^+^ population was clearly distinct for the other two while VE-Cad^+^GFP^−^ and VE-Cad^+^GFP^+^ cells were located closer to each other on the PCA plot (Fig. 2B). VE-Cad^+^GFP^−^ cells showed mainly high expression of endothelial genes (e.g. *Cdh5, Kdr, Tek, Pecam1*) but very little expression of hematopoietic and erythroid genes, while VE-Cad^−^GFP+ population displayed high expression of erythroid specific markers (e.g. *Gata1, Hbb-bh1, Epor*) and no expression of endothelial genes (Fig. 2C). For this reason, VE-Cad^+^GFP^−^ population was named Endothelial Cells (EC) and VE-Cad^−^GFP^+^ was denoted Erythroid (Ery). On the other hand, VE-Cad^+^GFP^+^ cells had high expression of both endothelial and erythroid genes (e.g. *Gata1, Epor* and *Hbb-bh1*) and low expression of other hematopoietic markers (e.g. *Itgam* and *Spi1*). The majority of the cells within each population showed homogenous expression of the majority of genes. Nonetheless, certain hematopoietic (e.g. *Itga2b, Itgab3, Ptprc* and *Gfi1b*) and erythroid specific genes (*Epor*) were more variably expressed in some of the populations. While the VE-Cad^+^GFP^+^ population had some *Spi1* (myeloid marker) expression, the majority of the cells did not express this gene (Fig. 2C). Likewise, *Runx1* was expressed in a small subset of VE-Cad^+^GFP^+^ (Fig. 2C). Among the 99 VE-Cad^+^GFP^+^ sorted cells, only a small subset (4 cells) expressed all together endothelial, hematopoietic and erythroid genes (Fig. S1). In summary, the VE-Cad^+^GFP^+^ population could represent a new hemogenic endothelium population that may directly give rise to erythroid cells: the Erythroid Hemogenic Endothelial Cell (Ery-HEC) population.

### Functional characterization of the Ery-HEC population

The function of the Ery-HEC population at E11.5 was evaluated by culturing these cells in the presence of OP9 stromal cells in a medium that supports the differentiation of hemogenic endothelium towards hematopoietic progenitors (Fig. 3A). The OP9 cell line was previously described to support the generation of hematopoietic progenitors from ES cells and hemogenic endothelium (Nakano et al., 1994; Swiers et al., 2013). To enable the growth of erythroid cells, we added erythropoietin (EPO) in the media. 500 Ery-HEC (VE-Cad^+^GFP^+^), 1,000 EC (VE-Cad+GFP^−^) or 1,000 Ery (VE-Cad^−^GFP^+^) cells from miR144/451^+/GFP^ E11.5 YS were FACS sorted on OP9 stromal cells. Following 4 days of culture in presence of EPO, the Ery population could not give rise to any colonies (Fig. 3B and 3C). However, EC were mostly generating white colonies (63% of the wells) and less often both white and green colonies (37% of the wells), which could be the progeny of EMP (Fig. 3C). Indeed, in our sorting strategy, we have not excluded CD45^+^ cells, which are in the final phase of EHT in the yolk sac. As for the Ery-HEC population, we interestingly observed that the majority of cultures gave rise to GFP^+^ colonies (45%) while wells producing only GFP^−^ or a mix of GFP^+^ and GFP^−^ cells were observed less often (27.5% for each type of colonies). This would indicate that even though this population is capable of giving rise to erythroid cells, the generation of white blood cell types such as megakaryocytes is also possible (Fig. 3B). This is consistent with the observation that rare cells in the Ery-HEC population are expressing *Runx1* (Fig. 2 and Fig. S1), which could generate these white colonies.

**Figure 3:**
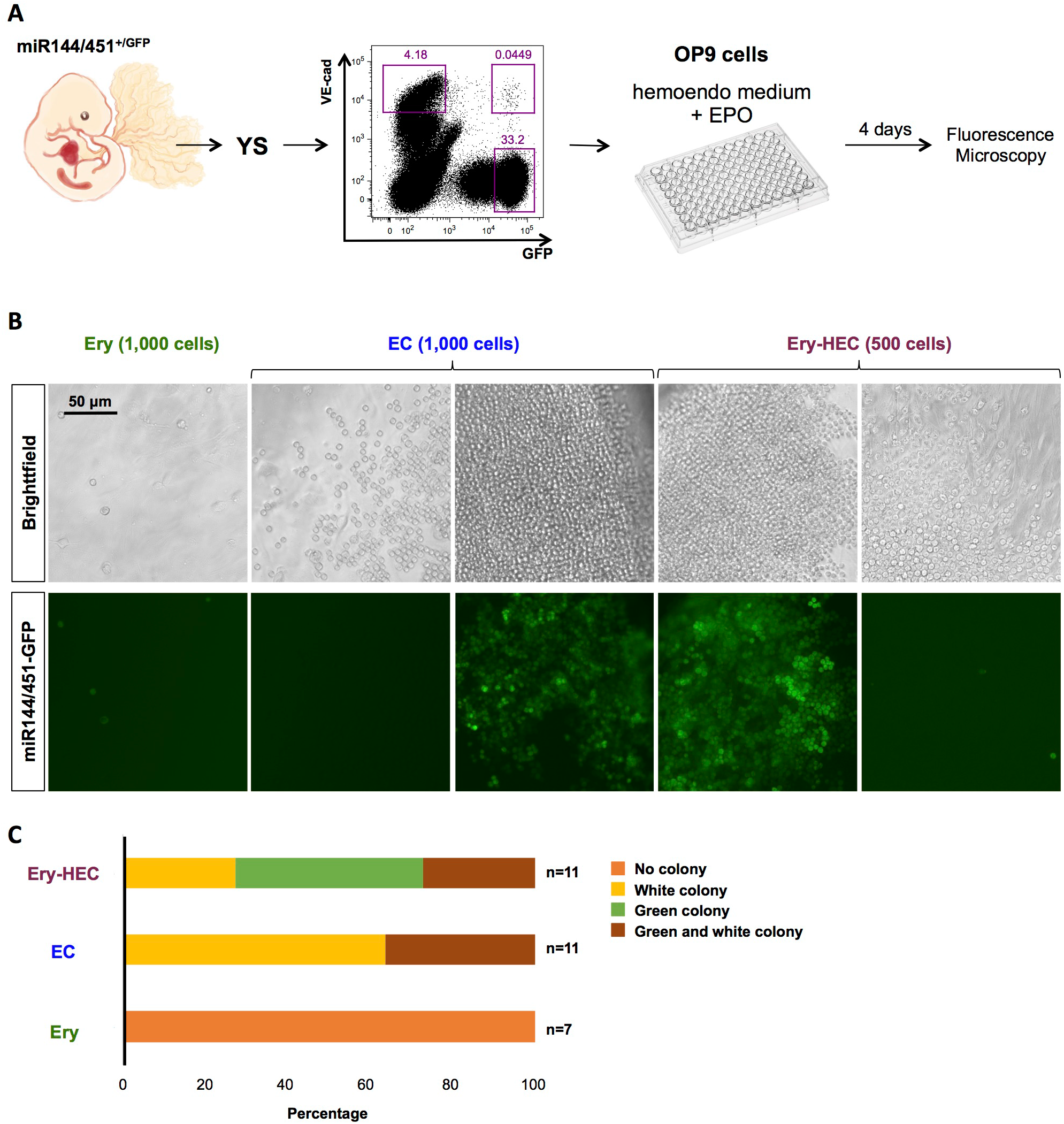
Ery-HEC can generate erythroid cells. (A) Schematic representation of the hematopoietic progenitor assay. 500 Ery-HEC (VE-Cad^+^GFP^+^), 1,000 EC (VE-Cad^+^GFP^−^) or 1,000 Ery (VE-Cad^−^GFP^+^) cells from miR144/451^+/GFP^ E11.5 YS were FACS sorted on OP9 stromal cells. The sorted populations are shown in the FACS plot in the purple squares. After 4 days in culture with an enriched hemogenic endothelium medium containing EPO, cells were analyzed by fluorescent microscopy. (B) Fluorescent and bright field images of wells seeded by Ery, EC and Ery-HEC populations after 4 days in culture. (C) Bar plots summarizing the percentage of wells with no colonies, white, green and mixed colonies. The number n of wells for each population is indicated.

### Runx1 is dispensable for the generation of Ery-HEC population

Since the single-cell q-RT-PCR results showed that the expression of Runx1 in the Ery-HEC population was very rare (Fig. 2C), we decided to thoroughly evaluate whether Runx1 is necessary for its generation. To achieve this, we generated Runx1^+/−^ animals (Fig. S2) and crossed them with miR144/451^+/GFP^ to obtain Runx1^+/−^ : miR144/451^+/GFP^ mice. Animals of this genotype were then crossed together to obtain Runx1^-/-^ : miR144^+/GFP^ and Runx1^-/-^ : miR144^GFP/GFP^ embryos. We then isolated YS from these embryos and sorted Ery-HEC by FACS for single-cell q-RT-PCR analysis (Fig. 4A). We were able to obtain 24 cells, with an expression profile highly similar to the wild type Ery-HEC cells (Fig. 4B and 4C), showing that Runx1 is dispensable for the generation of this population.

**Figure 4:**
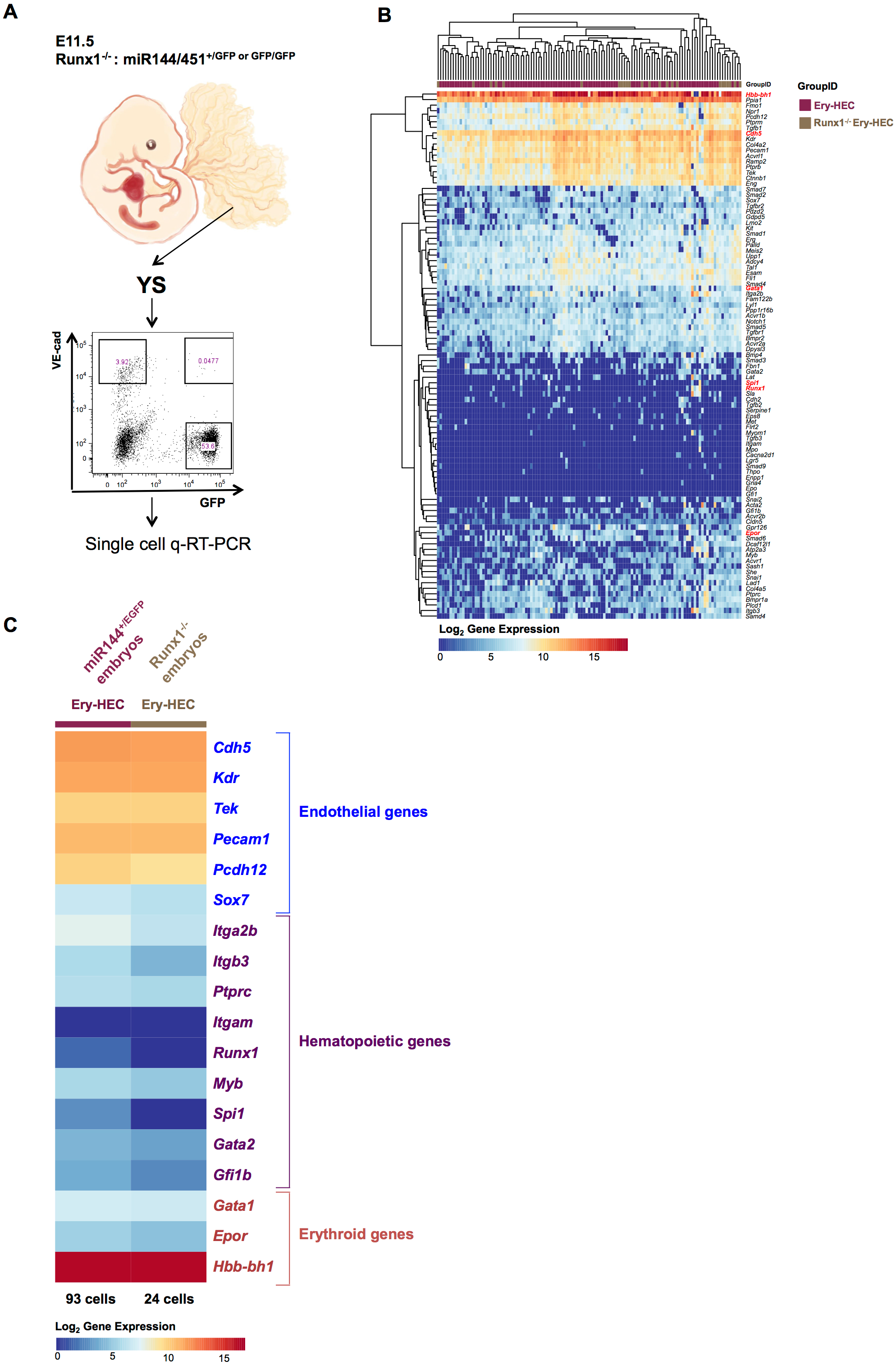
The Ery-HEC population is present in Runx1^-/-^ yolk sac. (A) Schematic representation of the experimental design for the isolation of VE-Cad^+^GFP^+^ cell population for single cell q-RT-PCR, using Runx1^-/-^ : miR144/451^+/GFP^ or Runx1^-/-^:miR144/451^GFP/GFP^ embryos. (B) Hierarchical clustering of wild type and Runx1^-/-^ Ery-HEC single cells. Red dots highlight a selection of genes, which are from top to bottom: *Hbb-bh1, Cdh5, Gata1, Spi1, Runx1* and *Epor*. (C) The average expression of each listed gene was calculated for the 2 indicated groups of cells. The figure shows an expression heatmap of several endothelial, hematopoietic and erythroid genes. The number of cells contained in each group is stated at the bottom of each column. See also Figure S2.

### The Ery-HEC population expresses genes characteristic of the primitive erythroid lineage

We have shown that the Ery-HEC population could be identified using the miR144/451^+/GFP^ mouse model together with the VE-Cad marker. The expression of *Hbb-bh1* suggests that these cells have a signature compatible with primitive erythropoiesis. However, it has been shown that this gene is also expressed by fetal definitive erythroid cells (Erythron data base-Kingsley et al. 2013). Therefore, to have a more complete understanding of the nature of these cells, we performed RNA sequencing. We sorted Ery-HEC cells from miR-144/451^+/GFP^ E11.5 YS and used EC and Ery populations as controls (Fig. 5A). We detected a total of 13,081 expressed genes (Supplementary File S4).

**Figure 5:**
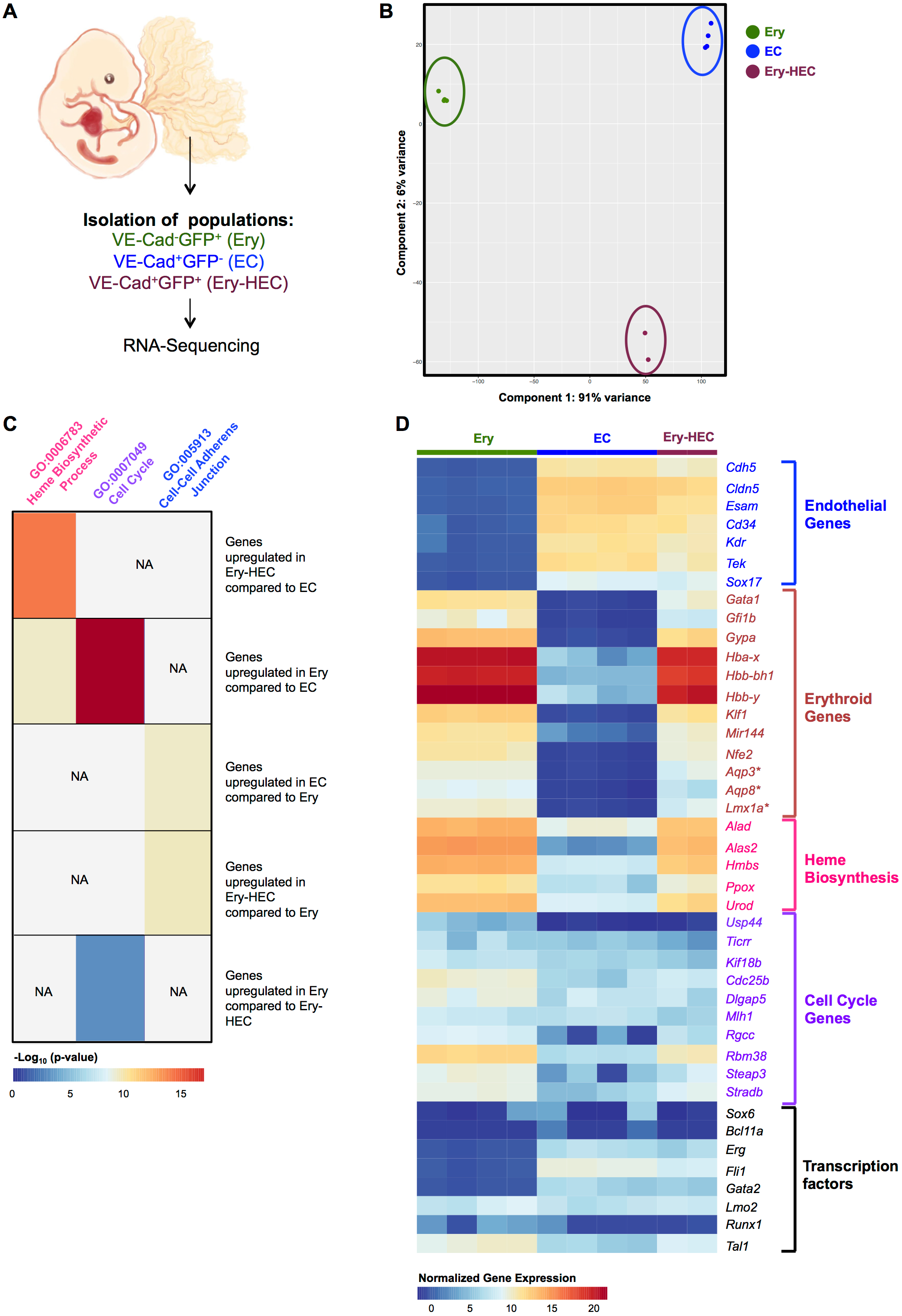
The Ery-HEC population expresses embryonic hemoglobins and markers of primitive erythropoiesis. (A) Schematic representation of the experimental protocol for bulk RNA sequencing of miR144/451^+/GFP^ YS VE-Cad^+^GFP^+^ (Ery-HEC) populations (1 sample of 5 and 1 sample of 11 cells), at E11.5. VE-Cad^−^ GFP^+^ (Ery) (4 replicates of 25 cells) and VE-Cad^+^GFP^−^ (EC) (4 replicates of 25 cells) cells were used as controls. (B) PCA plot showing the distribution of the different samples. (C) Heatmap summarizing the results of the GO term analysis with the DAVID Bioinformatics Resources. The analysis was performed on the statistically significant genes derived from each pairwise comparison. The heatmap displays the −log_10_(p-values) for each indicated GO term and group of genes. The p-values were calculated using the FDR method. NA means “not applicable” because the indicated genes were not enriched for the particular set of genes (i.d. there were no available p-values). (D) Expression heatmap of selected genes. Gene names followed by “*” indicate primitive erythroid genes. See also Table S2 and Supplementary File S4.

The PCA showed that the different sample groups have very distinct expression profiles (Fig. 5B). There were 4,115 differentially expressed genes (|log_2_ fold change|>1, p-value<0.01) between EC vs Ery, 2,188 between Ery-HEC vs Ery and 513 between Ery-HEC vs EC. Using these gene lists, we performed Gene Ontology (GO) analysis (Fig. 5C, Supplementary File S4 and Table S2).

The Cell-Cell Adherens Junction GO term (GO:005913) was more enriched for the two VE-Cad^+^ populations compared to the Ery population consistent with the necessity of endothelial cells to establish physical links to adhere to each other. Moreover, EC and Ery-HEC are clearly expressing endothelial genes such as *Cdh5* and *Cldn5* (Fig. 5D). The genes coding for the key transcription factors Erg, Fli1, Gata2, Lmo2, Runx1 and Tal1 were all expressed in the endothelial populations when only Lmo2, Tal1 and Runx1 could be detected in the erythroid subgroup (Fig. 5D). In line with the erythroid phenotype of the Ery population, there was a significant increase of genes associated with Heme Biosynthetic Process GO term (GO:0006783), e.g. *Alad* and *Alas2*, which is a crucial step in the production of hemoglobin. These genes were also significantly expressed in the Ery-HEC population supporting the idea that they could make hemoglobin. Interestingly, genes associated with cell cycle (GO:0007049) were more highly expressed in Ery population compared to the two endothelial populations suggesting a higher proliferative capacity. We also confirmed that Ery-HEC and Ery cells express miR144 which was used as a marker for both populations and lack Runx1 expression as shown before (Fig. 5D).

Finally, to know if the Ery-HEC cells are associated with the primitive erythroid program, we looked for specific primitive genes such as *Aqp3, Aqp8* and *Lmx1a* (Erythron data base - Kingsley et al. 2013) as well as embryonic hemoglobin genes (*Hba-x, Hbb-bh1* and *Hbb-y*). They were both expressed in the Ery-HEC and Ery populations. Interestingly, the adult hemoglobin *Hbb-b1* was not detected. Moreover, the genes coding for Bcl11a and Sox6, two transcription factors involved in the hemoglobin switching (Sankaran et al. 2009, Yi et al. 2006) were not expressed (Fig. 5D). Overall, this implies that the Ery-HEC population could be a source of primitive erythroid cells at much later stages than previously thought (Isern et al., 2011; Palis et al., 1999).

### Gata1 can induce the erythroid program in endothelial cells

The existence of a putative HE population specific to the erythroid lineage prompted us to ask how this program could be initiated in vascular cells. Our main candidate for this was *Gata1* because it has been described as a pioneer transcription factor to open silent chromatin. Indeed, ectopic expression of Gata1 in monocytes initiated erythroid and megakaryocyte gene expression (Kulessa et al., 1995; Visvader et al., 1992). In addition, *Gata1, Tal1, Lmo2* and *cMyc* have been shown to reprogram fibroblasts into embryonic erythroid progenitor (Capellera-Garcia et al., 2016). Since all endothelial cells express already *Tal1* and *Lmo2*, we reasoned that the induction of Gata1 expression might be enough to initiate the erythroid program in these cells.

To test this hypothesis, we used the Embryonic Stem Cell (ESC) differentiation into embryonic blood cells, a faithful model of yolk sac hematopoiesis. We first checked if endothelial cells co-expressing *Gata1* and *Hbb-bh1* could be produced from ESCs. We first differentiated the cells into Flk1^+^ mesodermal cells containing the *in vitro* equivalent of hemangioblast. These cells were later grown in the hemangioblast differentiation assay in the presence of IL6 and VEGF. After 1.5 days, VE-Cad^+^CD41^−^ endothelial cells were FACS-sorted and analyzed by single cell q-RT-PCR (Fig. S3). Out of 165 endothelial cells, 9 co-expressed *Gata1* and *Hbb-bh1* showing that these cells could also be produced *in vitro* from ES cells.

We next generated a doxycycline (dox) - inducible Gata1 ESC line (Fig. 6A) to test the role of Gata1 in the initiation of the erythroid program in endothelial cells. We demonstrated by western blot that the GATA1 protein was successfully expressed following dox treatment of ES cells (Fig. 6B). We next induced Gata1 expression during the hemangioblast differentiation assay and analyzed the phenotype of the cells after 1, 2 and 3 days of dox treatment. There was a significant reduction of endothelial cells frequency at all time points (Fig. 6C). There was also an increase of CD41^+^ frequency especially after one day of culture. The frequency of CD41^+^ cells was quite similar between the −dox and +dox conditions later on (Fig. 6C). To have a more detailed understanding of Gata1 expression impact in endothelial cells, we FACS-sorted VE-Cad^+^CD41^−^ cells from day 1.5 of hemangioblast differentiation assay. We also isolated VE-Cad^+^CD41^+^ cells, which are at a step further in the EHT and coexpress both endothelial and blood genes (Bergiers et al., 2018). These cells were cultured in hemogenic endothelium (HE) mix in absence or presence of dox for 48 hours (Fig. 6D). The q-RT-PCR analysis showed that the Gata1 over-expression led to the up-regulation of *Itgb3* (expressed by early progenitors and megakaryocytes), *Epor, Gfi1b, Hbb-y* (expressed by primitive and fetal erythrocytes) and *Hba-x* (specific to primitive erythroid cells) in the VE-Cad^+^CD41^+^ cell population suggesting the induction of the erythroid program (Fig. 6E). In contrast, the myeloid (*Spi1*) and endothelial (*Cdh5* and *Kdr*) programs were down-regulated (Fig. 6E). The impact of Gata1 over-expression on VE-Cad^+^CD41^−^ cells was less strong than on the other population. However, we observed a similar up-regulation of *Itgb3* and *Gfi1b* genes (Fig. 6E). *Hbb-y* and *Hba-x* expression was also higher following Gata1 induction although the change was not statistically significant (Fig. 6E). This was mostly due to large variability between the biological replicates (Fig. S4). Finally, CFU-assays were done in absence of dox following the HE cultures and we found a consistent and significant increase in the number of erythroid colonies (between 2 and 4-fold) following Gata1 over-expression for both populations (Fig. 6F). These results show that Gata1 can indeed trigger the erythroid program in endothelial cells.

**Figure 6:**
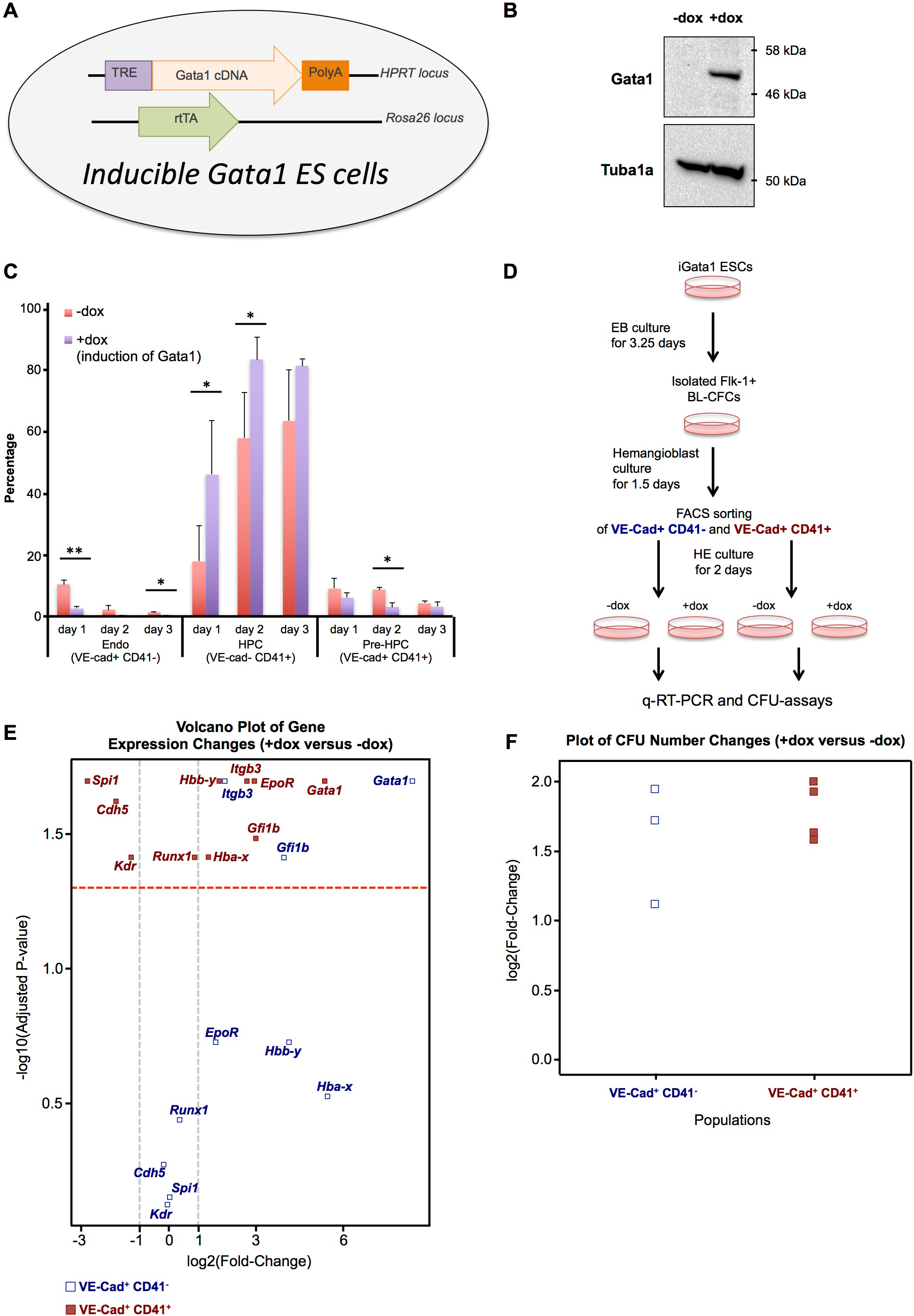
Over-expression of Gata1 induced the erythroid program in endothelial cells. (A) Scheme describing the iGata1 ESC line. (B) The western blot shows the protein expression of Gata1 following a 24h dox treatment of iGata1 ESCs. The loading control Tubulin Alpha 1a (Tuba1a) was used. (C) Average percentage of the indicated populations after 1, 2 and 3 days of culture with and without dox. Error bars are standard deviation. N=3. (D) VE-Cad^+^CD41^−^ and VE-Cad^+^CD41^+^ cells were sorted from day 1.5 iGata1 hemangioblast culture and seeded in hemogenic endothelium culture with and without dox. q-RT-PCR and CFU-assays were performed. (E) Volcano plot showing the log2(Fold-Change) values between +dox and −dox conditions following the q-RT-PCR of the indicated genes. The two grey dashed lines cross the x-axis at log_2_(Fold-Change) of −1 and +1. All −log_10_(Adjusted P-value) values above the red dashed line indicate statistically significant values. from the cells shown in D. N=3 (biological replicates). See also Supplementary Figure 3. (F) The scatter plot indicates the log2(Fold-Change) values between +dox and −dox conditions following the CFU-assays for the two indicated cell subsets. All the values are statistically significant (p-value <0.05, N=3 for VE-Cad^+^CD41^−^ and N=4 for VE-Cad^+^CD41^+^). See also Fig. S4.

## Discussion

In this study we have used single cell gene expression analysis to identify and characterize a new and rare population, Ery-HEC, with both endothelial and erythroid characteristics. Chen and colleagues previously suggested the existence of different types of hemogenic endothelium for EMPs and HSCs (Chen et al., 2011). However, this is the first time an endothelial population with apparent restricted primitive erythroid potential is found in the vasculature at such a late embryonic stage.

This was also the first time that this type of population was detected after the onset of both the primitive and EMP generating erythroid waves. While the first erythroid progenitors (EryP-CFC) are found in the Yolk Sac as late as E9.0, EMPs start to emerge at E8.25 from the newly formed vasculature (Frame et al., 2015; Palis et al., 1999). This new Ery-HEC population was observed in the YS from E9.5, increasing in numbers until E12, concomitant with primitive erythroid enucleation and the release of the first definitive erythroid cells derived from EMP, into circulation by the fetal liver (Kingsley et al., 2004).

We further showed that the Ery-HEC population expresses genes specific to the primitive erythroid lineage (e.g. *Aqp3, Aqp8* and *Lmx1a)*, (Kingsley et al., 2013). In addition, we showed that similarly to the first erythroid progenitors, Ery-HEC population could emerge in the absence of Runx1. Furthermore, Ery-HEC cells gave rise to either erythroid-cells (miR144/451^−^GFP^+^) *ex vivo* but also white blood colonies. Even though miR144/451 is expressed mainly in erythroid cells, it has also been shown to be present in a small fraction of CMP and PreMegE (Rasmussen and O’Carroll, 2011). Moreover, a small fraction of the Ery-HEC population expressed *Runx1*. Consequently, the Ery-HEC population might not exclusively give rise to primitive erythroid cells but could also be a source megakaryocytic progenitors. This could explain the white colonies observed in our OP9 culture (Fig. 3).

Even though the number of Ery-HEC cells found *in vivo* is quite low, these cells could later proliferate after losing their endothelial properties and become erythroid progenitors. Indeed we found that Erythroid cells had a higher expression level of cell-cycle genes compared to endothelial populations (Fig. 5).

Direct reprogramming of fibroblasts towards cells expressing primitive erythroid genes, has been recently shown, using the minimal combination of the transcription factors Gata1, Lmo2, Tal1 and c-Myc (Capellera-Garcia et al., 2016). Here, we demonstrate that the erythroid transcriptional program (both primitive and definitive as suggested by *Hba-x* and *Hbb-y* upregulation) could be activated in endothelial cells by overexpressing Gata1 specifically.

In conclusion, this newly described hemogenic population could constitute a novel pathway for the generation of erythroid cells and megakaryocytic progenitors at later stages in mouse development. The early phase of specification of this hemogenic population requires Gata1 but not Runx1. However, the role of Runx1 later in the production of erythroid and megakaryocytes remains unclear (Fig. 7).

**Figure 7:**
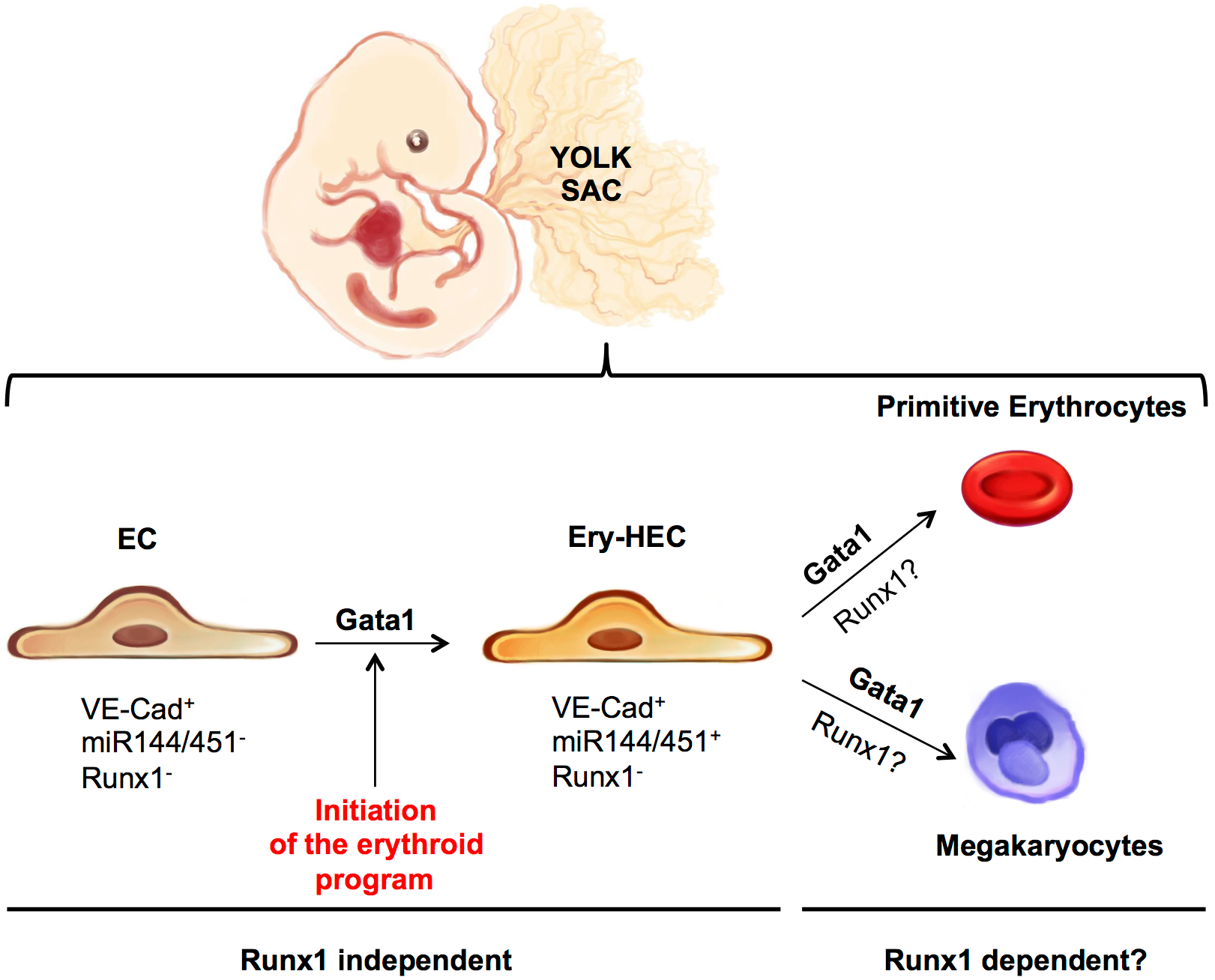
Model. The scheme describes the model derived from the results of the experiments shown in this paper. Gata1 is crucial to initiate the erythroid program and the expression of miR144/451 microRNAs and remains expressed throughout the differentiation. The resulting Ery-HEC population would produce erythroid cells and possibly megakaryocytes. Runx1 is not necessary for the generation of Ery-HEC but its function for the later phases of differentiation remains unclear.

Finally, the fact that Gata1 overexpression could induce the erythroid program in endothelial cells without using Myc, an oncogene, could be applied in medical applications aiming at producing red blood cells from endothelial cells instead of fibroblasts.

## Materials and Methods

### Mouse lines, generation of Runx1 null allele mice and embryo dissection

The miR144/451^+/GFP^ mice were kindly provided by Dr. Dónal O’Carroll (University of Edinburgh) and were previously described elsewhere (Rasmussen et al., 2011) while Runx1^+/−^ mice were generated as followed: The Runx1 gene targeting strategy allows Cre-mediated deletion of exon 2. This deletion creates a translation frame-shift producing a truncated Runx1 protein lacking the DNA binding and transactivation domains. The Runx1 gene-targeting vector carries a neomycin resistance cassette flanked with two frt sites. The targeting construct was transfected in A9 (129/C57BL6 hybrid) ES cells. Following G418 treatment (ThermoFisher, Cat# 11811023), resistant ES clones were tested by Southern blot to identify the homologous genomic integration. Correctly targeted ES cells were then used to generate mice heterozygous for Runx1 floxed allele. These mice were then crossed to the X-linked FLP-expressing transgenic mice to remove the frt flanked Neomycin resistant cassette, and then to the X-linked Cre-expressing transgenic mice to remove exon 2 and create the Runx1^+/−^ mouse line.

For embryo retrieval, mice were mated together with C57BL6, overnight, and vaginal plugs were checked the next morning at E0.5. Pregnant mice were killed by cervical dislocation in the time-points between E9.0-E12.5 of gestation. Yolk Sac dissections were carried out as previously described (Müller et al., 1994). All experiments were performed in accordance with the guidelines and regulations defined by the European and Italian legislations (Directive 2010/63/EU and DLGS 26/2014, respectively). They apply to fetal forms of mammals as from the last third of their normal development (from day 14 of gestation in the mouse). They do not cover experiments done with day 12 mouse embryos. Therefore no experimental protocol or license was required for the performed experiments. Mice were bred and maintained at EMBL Rome Mouse Facility in accordance with European and Italian legislations (EU Directive 634/2010 and DLGS 26/2014, respectively).

### Embryo genotyping

Mouse embryos were genotyped using KAPA Mouse genotyping kit (KAPA Biosystems, KK7352). For each embryo the head was dissected and incubated at 75°C for 10min in an eppendorf tube with 10μl of 10x KAPA extract buffer, 2μl of KAPA Express Extract Enzyme and 88μl of deionized H_2_O, in a total volume of 100μl. After incubation, tubes were vortexed briefly and centrifuged at maximum speed for 1min. For the PCR reaction 1μl of the supernatant was used together with 12.5μl of 2X KAPA2G mix, genotyping primers (10μM) and brought up to a final volume of 25μl using deionized H_2_O. PCR was performed in accordance to the protocol suggested in the kit and a 2% agarose gel was used for electrophoresis.

The primers for miR144/451^+/GFP^ and Runx1^+/−^ genotyping are listed below:

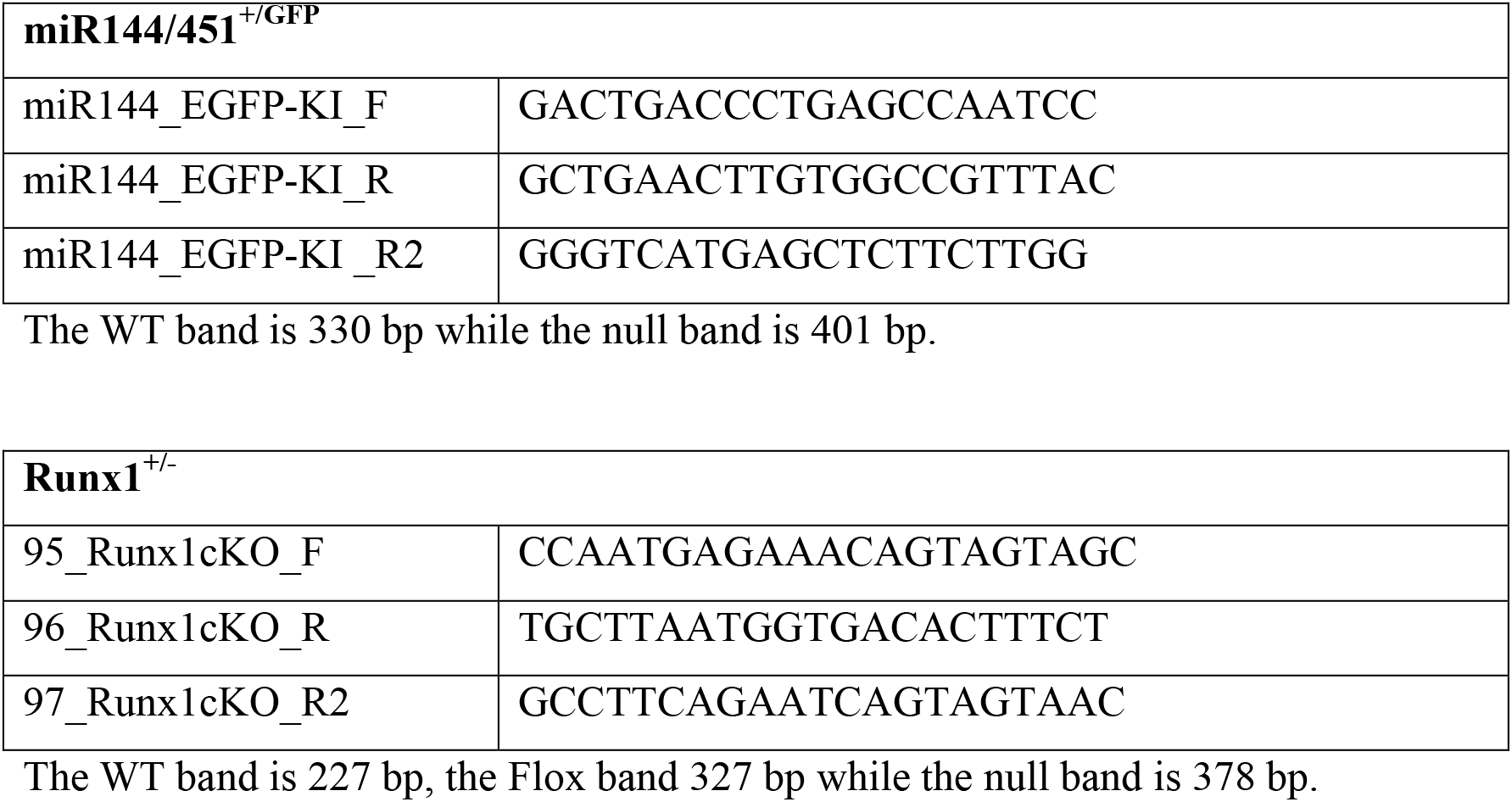

### Generation of inducible Gata1 ESC line

The inducible Gata1 (iGata1) ESC line was generated using the inducible cassette exchange method described previously (Iacovino et al., 2011; Vargel et al., 2016). The Gata1 coding sequence was tagged in 5’ with a Flag-tag, synthetized and cloned into a pUC57 Simple plasmid by the GenScript Gene Synthesis Service (http://www.genscript.com/gene_synthesis.html). The Gata1 coding sequence was then cloned from the pUC57 Simple into the p2lox plasmid (Iacovino et al. 2011). The subsequent p2lox plasmid was used to transfect A2lox.Cre ESC line (Iacovino et al. 2011) to generate the inducible Gata1 ESC line as previously described (Vargel et al., 2016).

### Western Blotting

Proteins were extracted from each sample (+Dox and - Dox) of iGata1 ES cells (approximately 6 million cells used) and 15 μg of each subjected to electrophoresis through a NuPAGE^®^ Tris-Acetate gel (Life technologies, cat. #LP0001) transferred onto nitrocellulose and probed with anti-Gata1 antibody N6 (Santa Cruz Biotech, cat. #sc-265) and an anti-rat secondary antibody (VWR, cat. #95058-824) before detection using chemi-luminescence.

### ES cell culture

Runx1+^/hCD4^ (Sroczynska et al. 2009) and iGata1 ES cell lines were maintained in DMEM-ES culture medium: DMEM (Dulbeco’s modified Eagle medium) -KO (Invitrogen, 10829018), 15% Foetal bovine serum (FBS) (PAA Clone, A15-02), 0.024μg/ml LIF (produced by the protein expression facility at EMBL, Heidelberg), 0.12mM β-Mercaptoethanol (Gibco^®^, 31350-010) and kept in an incubator at 37°C, 5% CO2 and 95% relative humidity (RH).

To improve cell adhesion, plates were coated with 0.1% gelatin (BDH, 440454B) in PBS (phosphate-buffered saline) for 20 minutes prior to cell seeding. TrypLE-Express (Gibco^®^, 12605-010) was used to detach cells for further collection or passage. DMEM-KO stock medium was always supplemented with 1% Pen/Strep (Penicillin/Streptomycin) (Gibco^®^, 15140122), 1% L-Glutamine (Gibco^®^, 25030024) and 1%of non-essential amino-acids (Gibco^®^, 11140035). All mediums used for cell culture and differentiation, were sterile filtered after preparation with 0.22μm vacuum driven filtration unit (Millipore SteriCup, SCGPU 01RE).

### EB differentiation, hemangioblast and hemogenic endothelium cultures

Prior to EB culture, cells were passed twice on gelatin, first with DMEM-ES and afterwards with IMDM-ES: IMDM (Iscove’s modified Dulbecco medium) (Lonza, BE12-726F) with 15% FBS, 0.024μg/ml LIF and 0.12mM β-Mercaptoethanol. IMDM stock medium was always supplemented with 1% Pen/Strep and 1% L-Glutamine. Cells were harvested and cultured in petri dishes at a final density of up to 0.3×10^6^ per dish, with EB medium: IMDM, 10% FBS, 0.6% Transferrin (Roche, 10652), 0.03% MTG (Sigma, M6145) and 50mg/μl Ascorbic Acid (Sigma, A4544).

After 3.25 days of culture, EBs were harvested for hemangioblast assay as described previously (Vargel et al. 2016). Briefly, Flk1+ cells from day 3.25 EBs were sorted using magnetic sorting (MACS^®^, Miltenyi Biotech) and plated in BL-CFC mix composed of IMDM, 10% FBS, 1% L-glutamine, 0.6% transferrin, 0.03% MTG, 0.5% ascorbic acid, 15% D4T supernatant, 0.05% VEGF (10μg/ml) (R&D, 293-VE-010), 0.1% IL-6 (10μg/ml) (R&D, 406-ML).

For hemogenic endothelium culture, FACS-sorted VE-Cad+CD41^−^ and VE-Cad+CD41+ cells were cultured on gelatinized plate at a density of 0.2 × 10^6^ cells per cm^2^ (cells isolated from day 1.5 BL-CFC culture). The media was composed of IMDM supplemented with 10% FBS (same as BL-CFC mix), 1% L-glutamine, 0.6% 30 mg/ml transferrin, 0.3% 0.15 M MTG, 0.5% 5 mg/ml ascorbic acid, 0.024% 100 μg/ml LIF (EMBL-Heidelberg), 0.5% 10 μg/ ml SCF (R&D systems, cat. #455-MC), and 0.1% 10 μg/ml Oncostatin M (R&D systems, cat. #495-MO).

### Hematopoietic Colony Forming Unit Assay

Cells from HE day2, previously treated with dox or not, were harvested and plated at a given density per dish (35×10mm, Corning) with 3ml of CFU medium. This medium contains IMDM + 15% PDS serum replacement (Antech), 10% PFMH-II (GIBCO, 12040-077), 1%L-Glutamine, 0.6% Transferrin, 0.05mg/ml Ascorbic acid, 0.03% MTG, 0.05μg/ml SCF (Peprotech, 52250-03), 0.025μg/ml IL-3 (R&D, 403ML), 0.025μg/ml GM-CSF (R&D, 425-ML), 0.005μg/ml IL-11(R&D, 418-ML), 0.02 μg/ml EPO (R&D, 959-ME-010), 0.01 μg/ml IL-6, 0.025μg/ml TPO (R&D, 488-TO-005), 0.005μg/ml MCSF (R&D, 416-ML-010) and 55% of 500g/L methylcellulose (VWR, 9004-67-5) which is added after the liquid components and the cells are mixed together. Three dishes (corresponding to 3 repeats) were prepared in total for each cell sample. Colonies were counted and averaged for each sample, based on their morphology (white, red and a mix of both mix) after 5 days of culture.

The cell counts from the CFU-assay experiment were analysed with a generalized linear model using quasi-Poisson distribution and logarithm as link function. One variable for the conditions (+dox and −dox) and a second variable to account for batch effects were used as predictor variables. In addition, the model included an offset to model rates instead of counts while accounting for differences in the total number of plated cells. The analysis was done separately for each of the two subsets of cells. The significance and fold-change of the contrast +dox versus -dox were reported.

### Immunostaining

Whole-mount immunostaining protocol was adapted from a previously described protocol (Corada et al., 2013). Whole 3D view of dissected embryo was performed using a Z1 light sheet microscope (Zeiss, Germany). Upon staining whole embryos were embedded in 2% low melting point Agarose (LMA, Promega) /PBS and placed at 4°C for agarose polymerization. Subsequently embryos were dissected sagittally, the embryo was scooped off and the 2 parts of the yolk sac were filled with 1% LMA and set at 4°C for polymerization. Each part was dissected in other 4 pieces, which were attached on a syringe tip and hanged up into the imaging chamber of the Lightsheet Z1. Solid-state laser exciting at 488 and 641 were used as illumination sources and 5X objective was used for imaging.

Upon imaging the dissected YS was immersed in PBS solution to allow agarose to solubilize, and mounted flat onto a glassslide for further analysis. Mosaic reconstruction was performed at 20X on a DSD2 spinning disk confocal (Andor) using LEDs exciting respectively at 480 and 640 nm. The entire montage was carefully analyzed in *xyz*, and putative double positive cells (VE-Cad^+^GFP^+^) were imaged at higher resolution in *xyz* to confirm their co-expression of GFP and VE-Cadherin 647 (clone BV13 unconjugated, eBiosciences, 14-1441-81).

### Flow cytometry and cell sorting

Yolk Sacs were disaggregated by adding 50μl of collagenase (Sigma, C9722-50MG) in 1ml of PBS 10%FBS and incubating for 30-40 min. at 37°C. To stop dissociation, 9ml of PBS 10%FBS was added after incubation. Cells were then stained with the appropriate amount of antibodies mix for 10 min. at room temperature, washed and filtered. For every 5 million cells, 1ml of antibody solution was used for staining. Cells were either kept in FACS medium (PBS 10%FBS) for FACS Analysis or IMDM 10%FBS for sorting. The viability staining dyes Sytox blue (Invitrogen, S34857) or 7-AAD (Sigma, A9400) were used to exclude dead cells and used at a final concentration of 0.1mM and 1mg/ml respectively.

Both FACS analysis and sorting were performed on FACS Canto and FACS Aria (BD Biosciences), respectively. The Flowjo software was used to analyze FACS data and all the FACS plots shown in the figures show live cell populations without doublets.

A list of the antibodies used in FACS experiments is described below:

| Antibody | Alternative name(s) | Clone | Fluorochrome | Catalogue number (eBiosciences) |
| --- | --- | --- | --- | --- |
| CD144 | VE-cadherin, CDH5 | BV13 | eFluor® 660 | 50-1441-82 |
| CD41 | Integrin-α2b (Itga2b) | MWReg30 | PE | 12-0411-82 |
| CD117 | c-Kit | 2B8 | APC | 17-1171-82 |

### Quantitative RT-PCR

RNA was extracted from inducible Gata1 ESCs using the miRNeasy Micro Kit (QIAGEN). Complementary DNA (cDNA) was synthesized with RevertAid H minus First Strand cDNA Synthesis Kit protocol (ThermoFisher Scientific) using random hexamer primer. The qPCR reaction was performed according to the KAPA SYBR Green Universal qPCR kit protocol, in the 7500 Real Time PCR System (Applied Biosystems). The following list of primers was used for KAPA SYBR Green Universal qPCR. Expression values were normalised to housekeeping gene *Ppia*.

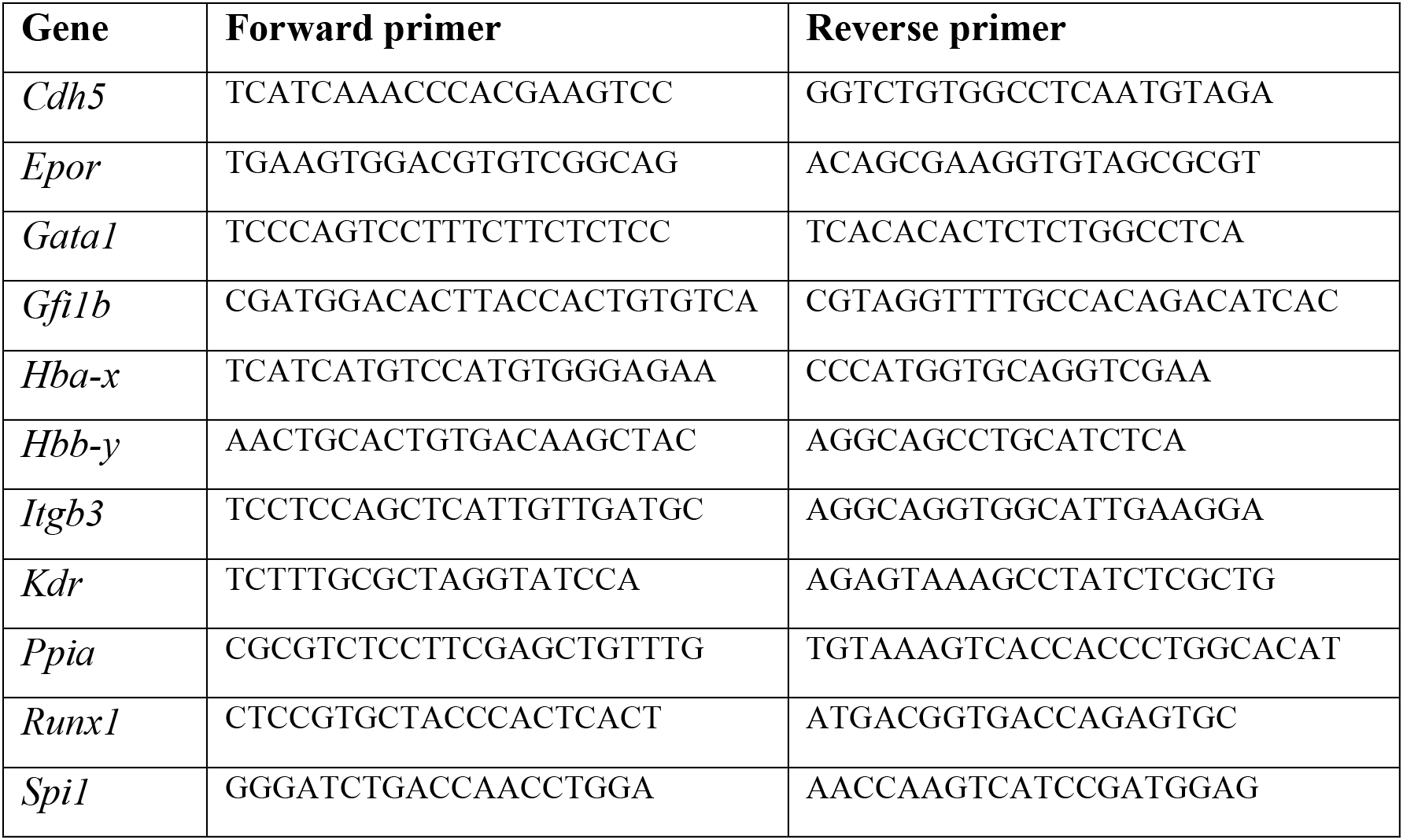

The relative RNA quantity was calculated the following way for each gene: CTtarget gene - CTreference gene (*P_pia_*) generating Delta CT values (Figure S4).

The results obtained from the q-RT-PCR as shown in Figure 6E were analysed with a 2-way ANOVA model accounting for the condition and the batch. The fold-change of the CT values and its significance between the +dox and -dox conditions was calculated for each gene separately. The Benjamini-Hochberg method was applied to adjust the p-values while correcting for multiple testing.

### Single cell q-RT-PCR

Single cell expression profiling was performed using a nested primer approach following the Fluidigm Advanced Development Protocol (section 41), in which 2 pairs of primers are used for each gene, one for reverse transcription and the other for qPCR. In brief, single cells from E10.5-12 YS and E11.5 AGM regions were FACS-sorted into 96-well plates (Bio-Rad Hard-Shell, HSP9611), containing 5μl of 2x reaction mix (Invitrogen, 11753) per well. The plates were shortly snap-frozen on dry ice after sorting and stored at −80°C.

Reverse transcription and nested-PCR were performed as previously described (Bergiers et al, 2018). The primers used for the 95 genes (Supplementary Table 1) are the same as in Bergiers et al (Bergiers et al, 2018).

### Hematopoietic progenitor Assay-OP9

Sorted cells from miR144/451^+/GFP^ E11.5 YS were co-cultured with OP9 stromal cells (OP9 ATCC^®^ CRL-2749TM) in 96 well plates with a modified version of hemogenic endothelium medium, containing IMDM 10% OP9 serum (ATCC^®^ 30-2020), 1% L-Glutamine, 0.5% Ascorbic Acid, 0.6% Transferrin, 0.03% MTG, 0.05mg/ml Ascorbic acid, 0.024μg/ml LIF, 0.05μg/ml SCF, 0.025μg/ml IL-3, 0.005μg/ml IL-11, 0.01μg/ml IL-6, 0.01 μg/ml OncostatinM, 0.001μg/ml bFGF (R&D, 233-FB) and 0.02 μg/ml EPO.

The OP9 stromal cells were plated the day before in 96 well plates previously treated with gelatin, at a final density of 3,000 cells per well and kept with OP9 medium. OP9 medium was replaced with the modified hemogenic endothelial medium before sorting.

### RNA sequencing

Cells from miR144/451^+/GFP^ E11.5 YS were FACS sorted into tube strips with lysis buffer containing 0.2% Triton X-100, oligo-dT Primer and dNTP mix and after snap frozen on dry-ice. Furthermore RT & template-switching, PCR preamplification (14 cycles for 25-cells and 16 cycles for 5-cells and 11-cells samples) and Nextera Library preparation were performed as described in the Smart Seq2 protocol (Picelli et al., 2014).

Sequencing data was analysed with the aid of EMBL Galaxy tools (galaxy.embl.de (Afgan et al., 2016)) for adaptor clipping (FASTX), mapping (RNA STAR) and obtaining raw gene expression counts (htseq-count). The R software (version 3.2.1, http://www.R-project.org.) was used to generate heatmaps and PCA plots by the DESeq2 package, and the web-based tools DAVID GO (https://david.ncifcrf.gov/tools.jsp) (Huang et al. 2009) was used to obtain lists of up- and-down regulated gene annotation clusters (See Supplementary File S4).

## Author Contributions

Irina Pinheiro, Conceptualization, Formal analysis, Supervision, Investigation, Visualization, Writing—original draft, Writing—review and editing; Özge Vargel Bölükbaşı, Conceptualization, Formal analysis, Investigation, Visualization, Writing—review and editing; Kerstin Ganter, Laura A. Sabou, Vick Key Tew, Giulia Bolasco, Investigation, Visualization, Writing—review and editing; Maya Shvarstman, Polina V. Pavlovich, Andreas Buness, Formal analysis, Visualization, Writing—review and editing; Christina Nikolakopoulou, Isabelle Bergiers, Investigation, Writing—review and editing; Valerie Kouskoff, Georges Lacaud, Conceptualization, Writing—review and editing; Christophe Lancrin, Conceptualization, Formal analysis, Supervision, Investigation, Visualization, Methodology, Writing—original draft, Project administration, Writing—review and editing.

## Acknowledgments

We thank Dr. Dónal O’Carroll (University of Edinburgh) for providing the miR144/451-GFP mouse line; Michael Kyba (University of Minnesota) for the A2.lox.Cre ESC line; Philip Hublitz (EMBL Genome Engineering Services) for designing the Runx1 targeting vector; Pedro Moreira (EMBL Transgenic Services) for Runx1 mutant ES cells and mice generation; Daniel Bilbao, Kalina Stantcheva and Cora Chadick (EMBL Rome FACS Facility) for cell sorting; Paul Collier, Bianka Baying and Vladimir Benes (EMBL Genomics Core Facility) for RNA-seq and single-cell q-RT-PCR; Catarina Pinheiro (Lisbon, Portugal) for scientific illustration. The European Molecular Biology Laboratory supported this work.

